# Machine Learning-Augmented Analysis of Nano-electrochemical Sensor Data for Predictive and Quantitative Assays of Complex Biological Samples

**DOI:** 10.1101/2025.11.27.691014

**Authors:** Chaitanya Gupta, Kishan Aryasomayajula, Sean R. Fischer, Zheng Jia, Nicki Chin, Hon Nguyen, Zohaib Baig, Jeremy Hui, Roger T. Howe, John M. Baldoni, Emmanuel Quevy

## Abstract

Analytical chemistry provides the content of nearly every scientific, technical and business decision relating to what atoms, molecules and devices are and how they interact in their environment. Various workflows come together in describing those interactions--separation, identification, quantitation, classification—to inform decisions across society. Each workflow has associated equipment, infrastructure and subject matter experts that integrate results for input into those decisions. However, those existing workflows are constrained by their evolution and often require integrated applications of the multiple analytical techniques integrated to form conclusions, thereby increasing cost, complexity and quality. This paper describes an alternative approach to addressing the many limitations of contemporary analytical chemistry. Here machine learning is leveraged to analyze high dimensional data derived from a nano-electrochemical sensor platform, to perform classification, identification and quantitation analyses in an integrative manner from a single-shot measurement. The approach described here for biological samples does not require sample-specific preparation or marker-specific probes and labels to identify and/or quantitate the analyte of interest, thereby liberating the physical measurement of the sample from chemicals and workflows that are tied to the underlying biochemical hypothesis. The nano-electrochemical sensor, as described in a prior publication, transduces the vibrational frequencies of multiple species in the sample into a collection of discrete electronic signatures that represent the sample in a high dimensional space, while using only 2-4µl sample volume. The sample analysis relies on training data to identify features in the high dimensional input dataset that are unique to the molecule or phenotype under consideration and distinct from the sample background matrix and/or a set of defined sample controls. The molecule- or phenotype-associated features can be tracked with clinically identified disease burden in the patient or with biomolecule concentration in the biological sample. The identification of patients with dementia from a plasma test is outlined here to demonstrate the applicability of this approach to classifying phenotypes associated with complex neurological disorders in a marker-agnostic manner. This approach is applied to a specific and quantitative assay of insulin Humalog and insulin Toujeo in a batch that comprises of a mixture of the two peptides, where Toujeo differs from Humalog in three amino acids residues only. We also describe the development of an assay for IL-6 in spiked human plasma samples with this method, thereby demonstrating the applicability of this analysis paradigm to more complex samples and protein-like species.

## Introduction

Analysis of samples from complex biological matrices is critical for disease diagnosis, mechanistic understanding of disease, chemical and biothreat characterization, drug discovery, and cell- and chemical-based manufacturing, amongst other processes. Traditional analyses rely on assay-specific reagents, instruments and workflows that tend to tailored to the hypotheses being tested, i.e. the investigator formulates a biological hypothesis pertaining to the sample that they wish to test, identifies the analytes that would best test the hypothesis and then leverages a suite of workflows to characterize and quantitate them in the sample [1]. We present a new analytical approach utilizing a simplified physical workflow, resulting in more information, and allowing one to address questions relating to classification (phenotype) molecular species (what entities contribute to the classification), and quantification (concentration of those molecules in the sample) in a single determination.

The method relies on the electrochemical excitation of the sample and measurement of the resulting high dimensional vibration frequency spectrum which contains a complex representation of the molecules within the sample. [2]. The map is analyzed to correlate with training set labels (healthy/disease; isoform A/isoform B; pass or fail specification; etc.) to create the determination model. The resulting classification model distinguishes the phenotypes. Test samples are analyzed with the determination model to determine the unknown sample phenotype. Using authentic standards with inferred association to the phenotype allows quantification of those molecules from the same sample spectrum by classical analytical workflows (e.g. standard curves). In summary, this approach creates a classification model with labeled authentic phenotypes, where prior phenotype associated knowledge allows for identification of the molecule species contributing to the classification and allows for quantification of those molecular entities using authentic samples as standards. In this paradigm, wet lab sample analyses are minimized by application of trained and validated models in a digital domain, rather than by the availability of different types of reagents, instrumentation, and workflow protocols in a wet lab environment.

Three separate applications are provided to demonstrate this new analytical capability.

1. A plasma-based model to differentiate age-matched patients with no clinical signs of dementia versus those that score high on the clinical dementia phenotype, based off the Clinical Demetia Rating (CDR) score.
2. Quantitation of long-acting (Toujeo/insulin glargine) and short-acting (Humalog/insulin lispro) in a mixture of the two pharmaceutical ingredients. These differ in composition by three amino acids at positionsB28, B29 and B30 of the
3. Assay for the inflammatory cytokine IL6 in human plasma samples, thereby demonstrating the ability to quantitate larger protein molecules in more complex matrices.

Measurements for the assays above require no more than 4µl of sample for measurement of the associated vibrational map. This sample volume enables analyses when sample availability is severely limited.

## Experimental Section

### (A) Electrochemical Sensor Preparation

The nanoscale electrochemical interface, at which the sample vibrational map is transduced, is realized as 50 nm diameter electrode that is located inside a 50 nm deep cavity from the sensing surface. The first reduction to practice for this sensor is where the exposed nanoscale electrode area is fabricated on the surface of a 250 µm diameter Pt (80%) – Ir (20%) STM tip substrate (Keysight Technologies, Santa Rosa, CA) that is coated with a 40nm HfO_2_ film using an atomic layer deposition (ALD) tool (Veeco, Cambridge, MA). A dual beam Hitachi 325 SEM/FIB system (EAG/Eurofins, Sunnyvale, CA) is used to mill the 50nm diameter hole in the HfO_2_ film to expose the underlying Pt-Ir electrode surface, which is subsequently functionalized with alkanethiol molecules with differing functional groups utilizing a 48-hour soak in an ethanolic bath. The ion-milled, functionalized tip substrate is carefully threaded through a pre-milled hole in a 3cm × 3cm × 1cm PDMS (Sylgard Dow Corning, Midland, MI) block, to ensure that 1µm of the STM tip substrate is exposed above the PDMS surface. Once loaded in the polymer gasket, the tip is connected to a flexible wire for electrical addressability by downstream custom instrumentation described below. The tip-gasket-wire assembly is loaded in the bottom half of a Teflon electrochemical cell, to which the top half is secured via screws. A compressed O-ring provides a leak-free seal between the top and bottom halves of the cell. Besides the FIB milled, PDMS mounted Pt-Ir working electrode, a 2 mm-thick gold wire (Alfa Aesar, Ward Hill, MA) and an off-shelf Ag/AgCl reference electrode (BASi, West Lafayette, IN) are also utilized to complete the traditional three-electrode electrochemical cell. Additional details on the sensor setup are available in a prior publication [2].

The subsequent reduction to practice for this sensing device, for manufacturability at scale, involves the fabrication of the 50 nm electrode in the 50 nm shallow cavity on a silicon wafer using a combination of custom 8-mask lithography, plasm oxide etching and Platinum-Aluminum-Titanium Nitride (Pt-Al-TiN) deposition processes (8-inch CMOS foundry, California). Metal layers are deposited by evaporation and patterned into thin leads by standard lithography and etching methods. Deep Ultraviolet (DUV) lithography and etching are used to define a 150nm hole in a Chemical Vapor Deposition (CVD)-grown silicon oxide layer on the wafer, in order to expose the underlying Pt metallic lead. The hole is filled by high dielectric constant hafnia (HfO2) deposited by conformal ALD, and the walls of the cavity and the coated surface of the Pt lead are selectively etched back by a directional plasma etching process to create the 50 nm cavity with the exposed Pt electrode lead. The nanoscale cavity is registered within a 10µm porthole defined by a patterned polyimide layer deposited on the CVD-grown oxide. The sensing device, as fabricated, is diced from an 8-inch wafer into ∼10k millimeter squared silicon dies. Several of these dies are laminated and glued on a flexible polymeric substrate. Millimeter-scale silver electrodes screen printed on the flexible substrate enable electrical connection between the nanometer-to-micrometer scale electrodes on the silicon die, when the silicon die is glue pressed onto the flexible substrate using an Anisotropic Conductive epoxy Paste (ACP)-mediated die attach process, where arrays of laser-milled porthole hundreds of microns in diameter on the flexible substrate are registered with the corresponding polyimide portholes on the silicon dies. The sheets of die-flex assemblies are cleaned by oxygen plasma to remove residual protection layers on the Pt surface and laser singulated into individual devices that are arrayed onto a tray. A custom molding process is leveraged to mold a cylindrical vial out of plastic, wherein the vial bottom has a millimeter size porthole and two gold-coated pins that are press fit into the molded vial while still hot. Glue is precision dispensed around the vial bottom porthole and the die-flex devices are picked and placed from their holding trays to the bottom of the plastic vial, by registering the porthole of the flex device to the vial porthole using computer vision. In these composite electrochemical cells, the sensor electrode is the working electrode, and the two gold pins serve as the counter and quasi-reference electrodes respectively. Each cell is thoroughly cleaned and tested for electrochemical functionality before using.

The cell, as prepared, is filled with 2.5ml of aqueous electrolyte that comprises of a stoichiometric ratio of potassium mono- and di-hydrogen phosphate (Sigma Aldrich, St. Louis, MO) in DI water at a total concentration of 100 mM, such that pH is maintained at 7.5. The electrolyte also contains a potassium ferri-/ferrocyanide redox couple in a 1:1 molar ratio, at a total redox concentration of 2mM. Samples that are presented for analysis are pipetted directly into electrolyte volume within the electrochemical cell. A schematic of the sensor-embedded consumable is provided in Figure 1a.

**Figure 1:**
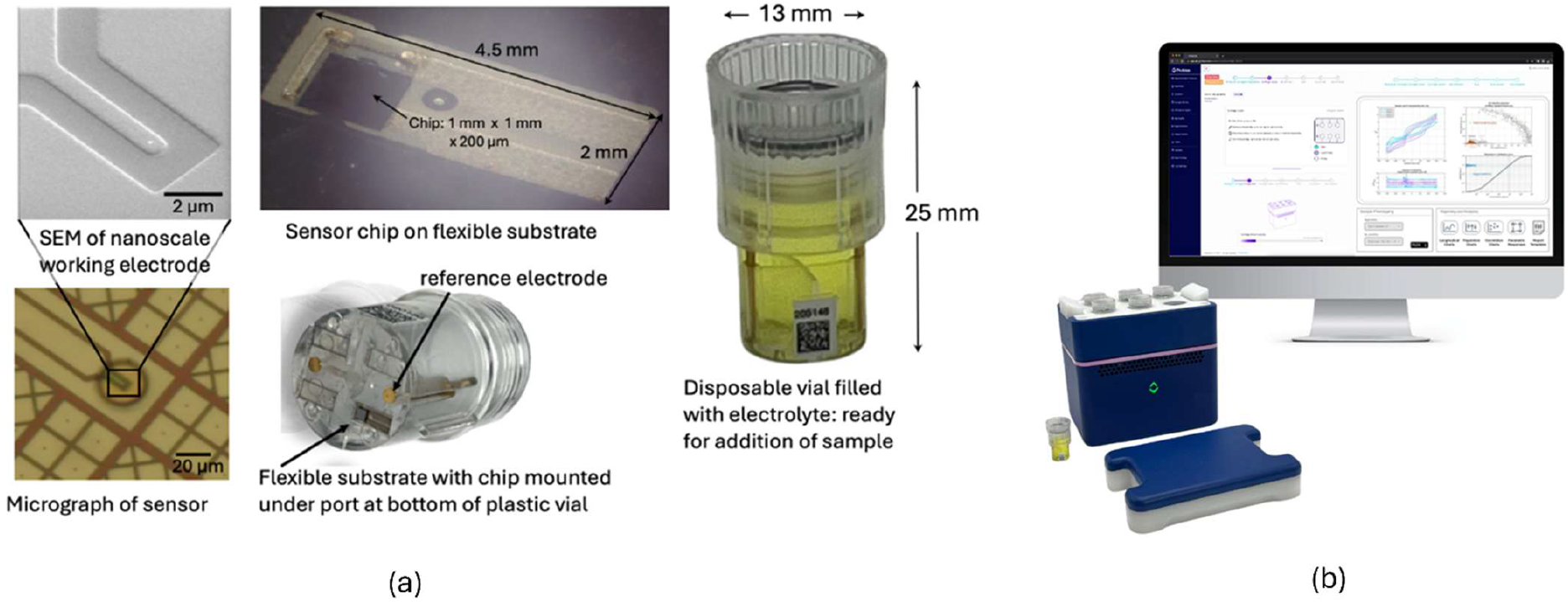
Schematics of (a) sensor ship, flexible substrate and consumable vial and (b) reader instrument on right that can measure six vials/samples simultaneously

### (B) Electrochemical Instrumentation and Measurement

The three electrodes of the electrochemical cell are connected to a custom, ultra-low noise potentiostat instrument, wherein the applied bias at the nanoscale electrochemical interface is cycled between -0.4V to 0.4V with respect to the Ag/AgCl reference electrode, or -0.2V to 0.7 V with respect to the quasi-reference gold electrode. The potentiostat is designed to apply bias to the interface with minimal contribution from the instrumentation electronic noise, while simultaneously recording the electrochemical charge transfer current with high resolution [4], which is critical to the measurement of the vibrational map [2]. A PC workstation hosting a custom MATLAB script automates the cyclical scan and data acquisition. Each electrochemical scan on a sample comprises of a thousand ensemble measurements of a) the voltage difference between the sensor node and the reference electrode and b) the resulting electrochemical charge transfer current across the electrochemical interface for a given voltage setpoint over a 1s data acquisition period, and the voltage setpoint is cycled between -0.4V to 0.4V, or -0.2V to 0.7V with 2mV increments. All samples are measured at least thrice, with six samples being measured simultaneously with the vial-based-reduction-to-practice for the sensor as depicted in Figure 1b. The complete scan dataset, acquired over a 30-minute period, comprises of the raw data that is automatically calibrated for offsets in the voltage and current measurement. The calibrated data is then filtered in the frequency domain by a MATLAB script that multiplies the Fourier transform of the signal with a bank of notch filters centered on frequencies of the observed electromagnetic interference aggressors in the experimental setup. The script subsequently performs an inverse Fourier transform to recover the filtered time-domain signal, with minimized contributions from the aggressors.

### (C) Sourcing and Handling Protocol for Biological Samples

Insulin glargine and insulin lispro used in the measurements described here were sourced by prescription (Walgreens, Sunnyvale, CA) and were certified to be 99.95% pure. These species were solubilized in phosphate buffer saline, and subsequently aliquoted into the aqueous electrolyte medium of the sensing cell. Multiple iterative dilutions of the aliquots were conducted to explore the sensitivity limits of the electrochemical detection method, while concurrently assessing specificity. In this paper, the peptide concentrations as present in the electrochemical cell range from 25ag/ml to 125ag/ml, with the two forms present in different ratios (0:1, 1:1, 1:2, 1:3) in each sample batch. Recombinant human IL6 (ProspecBio, E. Brunswick, NJ) was reconstituted in phosphate buffer saline and aliquots from the buffer mixture were used to spike the electrolytic buffer with the redox active species as well as human plasma, K2EDTA preserved, M/F pooled (BioIVT, Hicksville, NY) to simulate biological plasma samples with varying levels of inflammatory cytokine IL6. For demonstration purposes, we considered sixteen concentration values between 0.025pg/ml to 10ng/ml to explore the dynamic range of the measurement, consistent with expected variations in physiological IL6 expression [5]. Finally, 104 human plasma tubes were acquired from Boca Biolistics (Deerfield Beach, FL), which were obtained from patients that had been administered a comprehensive diagnostic assessment of clinical dementia that comprised of an MRI exam, subjective cognitive assessment, determination of clinical dementia rating (CDR) and mini-mental score examination (MMSE) as well as supporting assessments like family-, medical exam- and medication-histories. The patients in this study were equally split between those with diagnosed dementia (CDR > 0.5, MMSE < 24) versus patients with no cognitive impairment (CDR < 0.5, MMSE > 24). Both cohorts had an over-representation of women, with the gender ratio being 40:60, M:F in each, but were otherwise age and ethnicity matched. All plasma samples were thawed at most once before use in an ice bath and 4µl of the plasma specimens was pipetted into the electrochemical cell for measurement.

### (D) Data Analysis Methodologies

Data generated by an electrochemical scan on a sample is of the form of two matrices of size (801, 1000), where the measured current and voltage values at 801 set interface bias values are measured over 1000 times over the acquisition time of 1s. These matrices are subsequently post processed as described in (B) and reduced to a set of latent features using a group of “electrochemical learner functions” that are learned from the raw data. The measured electrochemical charge transfer current is represented as a weighted superposition of individual learner functions, where the charge transfer event is the outcome of several possible probabilistic paths, each path being denoted by the product of the weight and the learner function. The learner function itself measures the vibration-assisted, bias-dependent transition probability for electron transfer between nano-scale metal electrode and the redox-active species solubilized in the electrolyte, and the weight is representative of cumulative thermalized transitions that occur in conjunction with the bias dependent transfer [see Supporting Information]. We use an importance-sampling based approach to parameter estimation from a minimization of the least squares error between the raw data and weighted learner function representation, thereby regularizing the basis function representation of the charge transfer process [6]. The optimal span of the basis set is determined by minimizing the bias and in-set training error associated with the learner representation [see Supporting Information]. The end result of this transformation exercise on the current and voltage matrices is a feature set comprised of a down-sampled matrix of size (410, 300), where the values in the matrix define the unnormalized probability of the energy exchange interaction between an electron at bias value in the ordinate row and a vibrational mode at energy in the abscissa column.

The as-derived electrochemical feature set is flattened into a vector and subsequently input to one of multiple dimensionality reduction approaches that transform and/or reduce the input features into a feature that is the best representative of the target, where the target could be a sample phenotype or a concentration level associated with a specific molecular species. The dimensionality reduction approach is selected based on the end goal (classification/regression) of the analysis workflow as depicted in Figure 2. For triaging samples based on known phenotypes, an ensemble classifier approach comprised of a superposition of k-nearest neighbors (kNN), decision tree, random forest, support vector machine (SVM), linear, naïve Bayes and neural network classification models is utilized to associate specific features in the high dimensional datasets that correlate with binary label values in the training dataset. To minimize the possibility of overfitting, the classification algorithm preprocesses the input feature vector by normalization and feature selection, wherein the features are selected based on a) minimal variance, b) maximal correlation to label, c) maximal f-test score or d) maximal chi-square test statistic across phenotypic cohorts. The appropriate feature selection method is chosen based on classification accuracy from a simple SVM classifier that uses the feature-selected data. In addition, the overall weighted prediction assessment from the ensemble model is also regularized to reduce the possibility of overfitting [7]. The choice of component model parameters, weights for the ensemble prediction and the value of the regularization parameter are determined by hyper-parameter optimization across a validation dataset. For all classification tasks, a given measurement dataset is split 80:20 into an exclusive training + validation and test dataset, and the 80% dataset is further split into an exclusive 80:20 split between training and validation data. To explore the sensitivity of the trained model to the input, randomized splits of the training and validation data, this splitting process is also bootstrapped multiple times, where the model is trained on a disparate training + validation split and tested on a held-out test split, and the predictions from each bootstrap are assessed for concordance. Alternatively, the analysis workflow can also predict a concentration value for a specific species in a generated vibrational map of the sample, which is referred to as quantitative analysis in Figure 2. A similar ensemble-based approach is used for regression of concentration values of the analyte biomolecule in spiked biological matrices and in the blank electrolyte matrix, where the standard dilutions in the blank electrolyte matrix are prepared to emulate a ground truth representation of the vibrational spectrum of the analyte biomolecule free of matrix effects. The regression ensemble comprises of tree, random forest, SVM, kNN, linear, ridge and neural net regression models where the features are pre-processed like before, and the ensemble regularization, the component model parameters and the ensemble weights are hyper-parameter optimized on a randomly held out validation dataset. The training validation and test datasets are randomly and exclusively selected from the measurement datasets on the spiked samples, and the dataset splitting is also bootstrapped to ensure that the as-trained models are not overly sensitive to training data + validation data splits. The feature set generation and dimensionality reduction algorithms are scripted in MATLAB.

**Figure 2:**
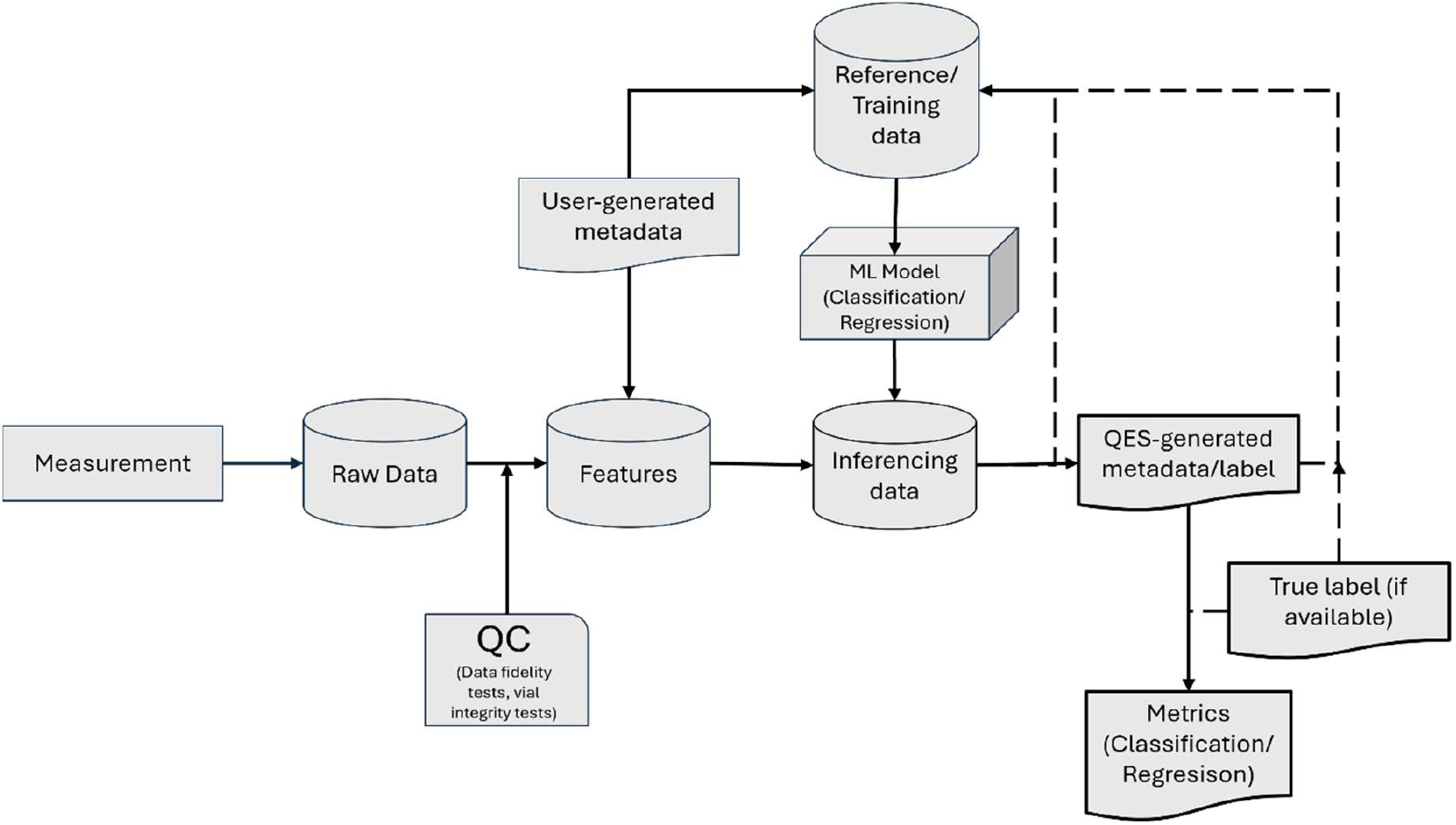
Schematic of data workflow in the analysis of sample measurements where the vibrations in the sample are measured electronically with a nano-electrochemical transducer. Raw data is screened and feature extracted using the procedure described in the Supplemental section. The sample feature representation can be inferred upon using an available model, for classification or quantitation (regression), and the labels generated from the inference process can be compared against the ground truth, if available, to define metrics for the digital assay. The measurement data can be subsumed into the training data to improve the machine learning model for future inferences.

## Discussion

### (A) Featurizing Electrochemical Data

In a prior publication [2], we have described a nanoscale electrochemical interface that generates a broad-spectrum vibrational fingerprint specific to a sample introduced into the electrolyte phase, when the low noise bias [4] across the interface is scanned across a range of values. This all-electronic transduction of vibrational frequency information occurs via a resonant exchange of charge between the metal and electrolyte phases, mediated by vibrational modes of species in the sample being analyzed. The exchange of energy between the transitioning electron and the intermediating vibrational mode can be resolved in the trace of the measured electrochemical flux if the transducing interface subscribes to a set of geometric-design and material-choice criteria [2], which enable the non-linear amplification of the electronic-vibrational interaction over the ambient thermal background and the simultaneous minimization of parasitic impedances like interface capacitance and/or mass-transported limited Warburg impedance. In this context, the measured electrochemical current can be represented as the probabilistic combination of a countably finite number of paths (*n*) for the transitioning electron, where each path includes resonant interaction with a specific vibrational mode as well as non-resonant thermalized activation processes that may precede or follow the resonant interaction:

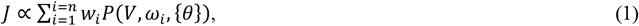

where *i* is an index over all transition channels, *w*_*i*_ represents the thermalized activation probability associated with channel *i* and *P*(*V, ω*_*i*_, {*θ*}) is the parametric probability of a vibrational frequency *ω*_*i*_ mediated resonant transition, and {*θ*} is the associated parameter set. A functional representation for *P*(*V, ω*_*i*_, {*θ*}) was presented in [2], specifically for the case when (a) the nanoscale interface is defined within a cylindrical 50nm diameter, 50nm deep cavity that acts as a spatial filter to exclude transverse polarization modes greater than 50nm in wavelength and when (b) the voltage noise energy spectral density associated with the applied bias is limited by the intrinsic noise power of the silver/silver-chloride reference electrode with respect with which the bias is applied. For these conditions, we have previously demonstrated that the electron transfer process is nonlinearly and resonantly coupled to vibrational states of interface polarization modes and as a result, characteristic signatures of the coupling can be observed in the measured charge transfer current. For purposes of subsequent analyses described in this paper, the measured current, as acquired from a traditional cyclic voltammetry experiment conducted on this nano electrochemical interface, can be decomposed into an equivalent basis set of functions, *P*(*V, ω*_*i*_, {*θ*}), where each basis function is weighted by *w*_*i*_. Figure 3a depicts a comparison between a typical experimental current-voltage trace acquired from a low voltage noise cyclic voltammogram on the nano electrochemical interface, as well as three different normalized current-voltage traces corresponding to three different resonant transfer channels, where each individual channel is associated with a particular vibrational frequency intermediating the transition. The individual channels depicted in Figure 3a correspond to individual basis functions *P*(*V, ω*_*i*_, {*θ*}) in the expansion shown in Equation (1).

**Figure 3:**
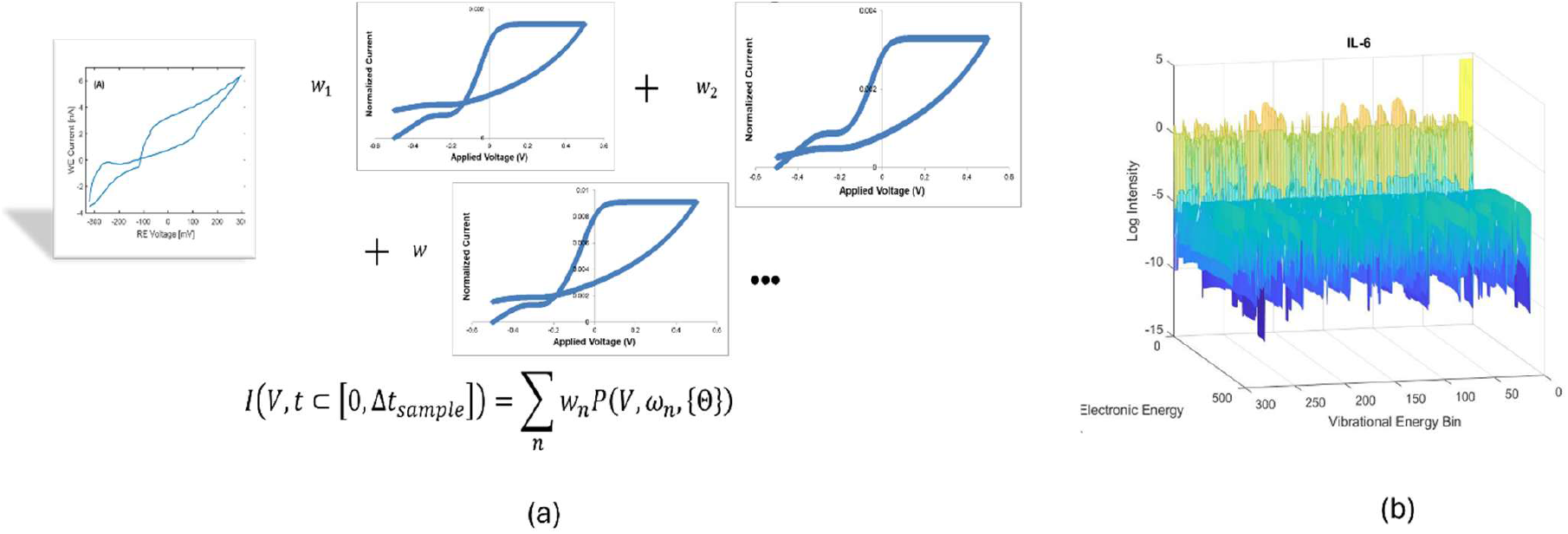
a) Current-voltage characteristics for a single charge transfer channel involving one contributing vibrational mode (*ω*_*n*_), where a weighted superposition of these individual characteristics enables the recreation of the measured transition flux at nano-electrochemical interface; b) equivalent spectral representation, after transformation of the raw current-voltage characteristic, of a sample of IL6 in PBS at 500pg/ml, from which a 4ul aliquot is measured in the electrochemical cell

A subset of the current-voltage-time data from the cyclic voltammogram experiment, randomly sampled without replacement, is used to estimate the parametric resonant probability basis functions. An importance-sampling Monte Carlo approach [6] is leveraged to estimate the unknown parameters by minimizing the MSE between the sampled, experimental current-voltage traces and the functional current-voltage traces generated from the parametric *P*(*V, ω*_*i*_, {*θ*}), as shown in Figure 3a. The weights or channel thermalized transition probabilities are determined subsequently from stochastic optimization of the least squares fit between the experimental current-time traces at different applied bias values, where the current-voltage-time data used to determine the weights are sampled from the data left out of the initial parameter estimation process. In summary, the parameter estimation enables a macroscopic description of the global current-voltage trace shape, and the weights represent a finer-grained characterization of the contribution of individual basis functions to the local trace. The estimates of the basis function parameters and the weights are performed for variable values of number of channels, *n*, for a population of current-voltage-time data acquired by scanning an ensemble of 50-100 samples. The optimal value for *n* is determined from the analytically calculated prediction-bias, prediction-variance tradeoff, as discussed in the Supporting Information section, and once optimal *n* is determined, it is fixed for all subsequent measurements performed using the same hardware (sensor and instrument combination). The set of parameters for the electrochemical probabilistic basis functions as well as the weights/thermalized transition probabilities constitute a feature-set that is leveraged for subsequent downstream analyses.

### (B) Feature Analysis Workflow

An example representation of transformed features generated from the featurization process in (A) is shown in Figure 3b, specifically for IL6 in electrolyte at a concentration of 500pg/ml in PBS, from which a 4ul aliquot is pipetted into the electrochemical cell for measurement. The x and y axes represent the electronic energy (in eV) and responding vibrational frequency bins (in eV), and the z axes measures the response intensity on a log scale. Once generated, the features are transformed through a sequence of nonlinear or linear transforms described here, to associate the sample vibrational features to (a) a pre-determined phenotype or (b) to a specific analyte in the sample and to determine the expression of that analyte in the sample matrix. A set of reference samples are leveraged to measure and generate the “ground-truth” feature representation of the phenotype or the analyte that is desired to be identified. For sample phenotype identification, a population of reference samples that are biochemically representative of the test samples are catalogued by metadata that informs the phenotype, where the phenotype could be determined by the environmental/situational context in which the sample was extracted or from an analogous gold standard assay that is used to assess the phenotype before the sample is made available for measurement with the nano electrochemical interface. In the case when we need to quantitate an analyte in a sample matrix, the reference population comprises of samples where the analyte is spiked in a blank matrix as well as in the relevant biological matrix, across an application-relevant dynamic range of concentration values and in increments that are small enough to provide the desired resolution in the concentration measurement. The reference samples are pipetted directly into the electrochemical cell and scanned to generate the raw current-voltage-time data, that is then subsequently transformed into electrochemical features.

Table 1 lists the many analyses that have been performed using the analytical approach that applies machine learning to the measurement data acquired from vibrational signal transduction at the nano electrochemical interface. In this paper, we describe three specific experiments in detail and the other assays will be described in follow-up publications. First, we triaged 104 plasma samples extracted from patients for whom clinical dementia was assessed using a combination of diagnostic MRI scans, subjective cognitive assessments, determination of clinical dementia rating (CDR) and mini-mental score examination (MMSE). The blood from these patients was spun down into plasma and stored with EDTA preservative. The plasma samples were frozen at -80C after preparation and were thawed only once before use. 4µl aliquots from each specimen was pipetted into the electrochemical cell for measurement by the sensor. The measurements of the plasma samples were made with the vial-based reduction-to-practice for the sensor. Every specimen was scanned three times per sensor, where each scan cycles from -0.2V to 0.7V and back on the applied bias scale, and measurements of the specimen were repeated at least six times with six different sensors. The applied voltage is referenced with respect to the quasi-reference gold coated electrode pin in the vial. Each scan generates a thousand steady-state instances of the measured electrochemical current crossing the nano electrochemical interface and the measured difference between the reference and nanoscale working electrodes. The current and voltage power spectral density, averaged over the applied bias values, are estimated from these measurements and banks of notched frequency filters are applied to the raw data to remove electromagnetic aggressors that may manifest in the measurements. The averaged power spectral density transforms are also monitored against expected baseline values, as determined from expected performance of instrument readout circuitry, to screen out measurements that are divergent from the baseline. The post-processed data is pipelined through the featurization process to generate electrochemical basis functions and weights for downstream analysis – the optimal ***n*** value is set to 300 from determination of optimal bias-variance tradeoff, as estimated from analytical measures of bias and variance in the electrochemical basis function representation. The as-determined electrochemical features are then input into a sequence of linear and/or nonlinear transforms designed to correlate specific properties of the sample, as characterized by the sample metadata/labels to the featurized data. The full measurement dataset on the human plasma samples is split 64:16:20 into training, validation and test data for analysis, where each cohort (control, dementia) is equally represented. For phenotype identification of dementia in the human plasma samples from controls, we use an ensemble model prediction approach, where the ensemble comprises of the following types of classification models: SVM, linear, neural network, kNN, decision tree, random forest and naïve Bayes. The ensemble model works on a pre-selected feature dataset, where the relevant features are selected either based on generalized statistical properties, like minimal variance within phenotypic cohorts, or based on statistical association to the phenotypic label. A simple model, namely a classification SVM, is used to determine the strength of association of features to the labels. The individual model parametrization, ensemble weights and ensemble regularization are hyperparameter optimized over a subset of the training data that is earmarked for validation. The optimized model is used to infer the labels on the held-out test dataset, and the accuracy of the as-trained classifier can be determined by assessing concordance to the actual label values. The receiver operator characteristic (ROC) for the classifier is graphed in Figure 4a, for 100 bootstraps of the training, validation, test data split – each bootstrap results in a classifier that is 100% sensitive and 100% specific for the corresponding test data, indicating that the classification approach is independent of the input training + validation dataset used to train the specific classifier used in that bootstrap iteration. We repeat the classification exercise on the same sample dataset, but with the patient gender as the data label in Figure 4b. Again, for bootstrapped training, validation and test splits, the classifier exhibits perfect accuracy in identifying male from female patients for this cohort. Notably, the new gender-specific classifier was re-trained and tested on the same patient cohort using different metadata associated with the sample, without requiring any new sample measurements. We expect the high classification accuracy depicted in Figure 4 to be on account of the small sample size of the sample cohort, despite the 20% leave-out strategy utilized for testing samples that are blinded with respect to the model trained within a specific bootstrap iteration.

**Table 1:**
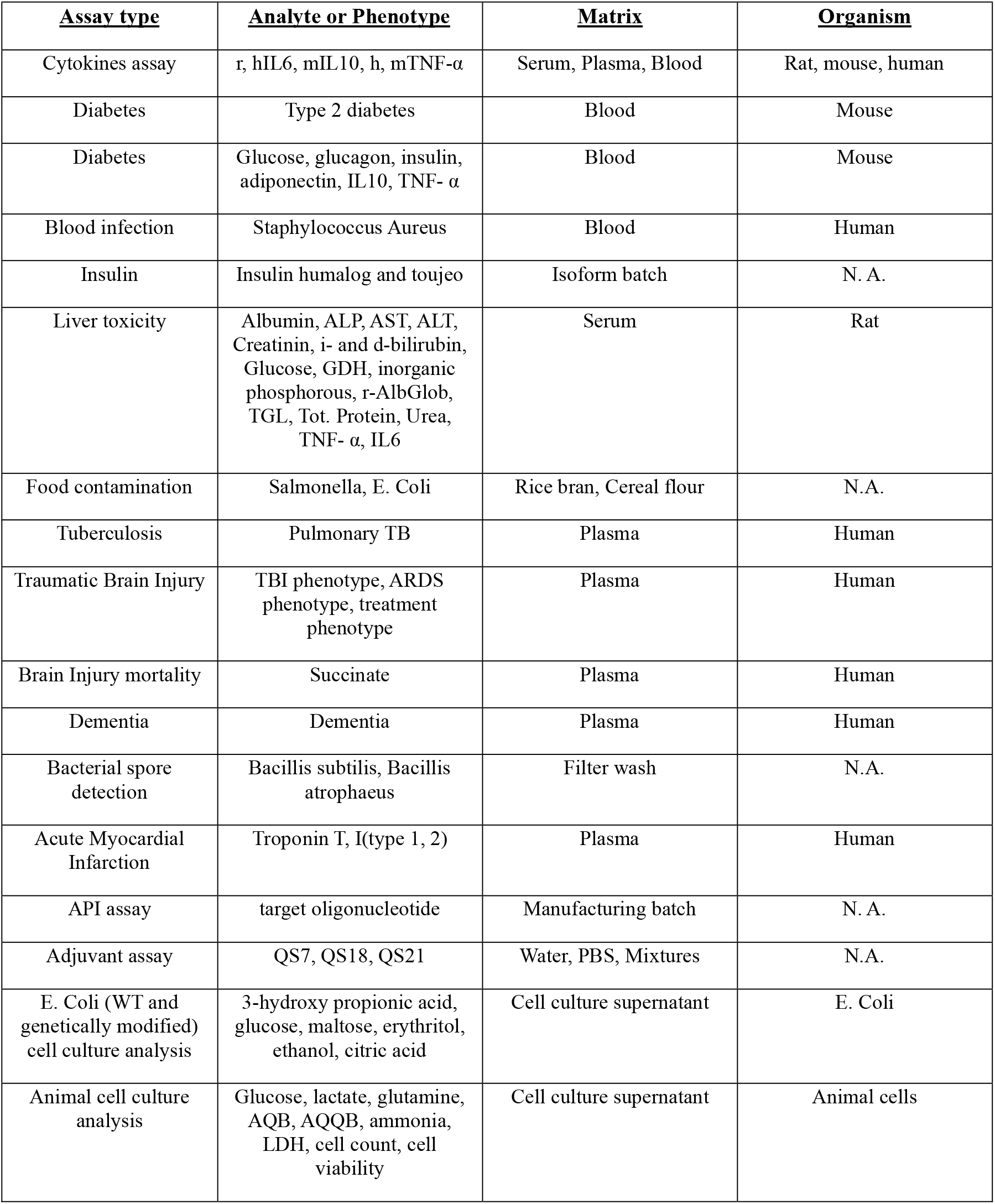
examples of analyses performed using machine learning applied to vibrational measurements from the nano electrochemical transducer.

**Figure 4:**
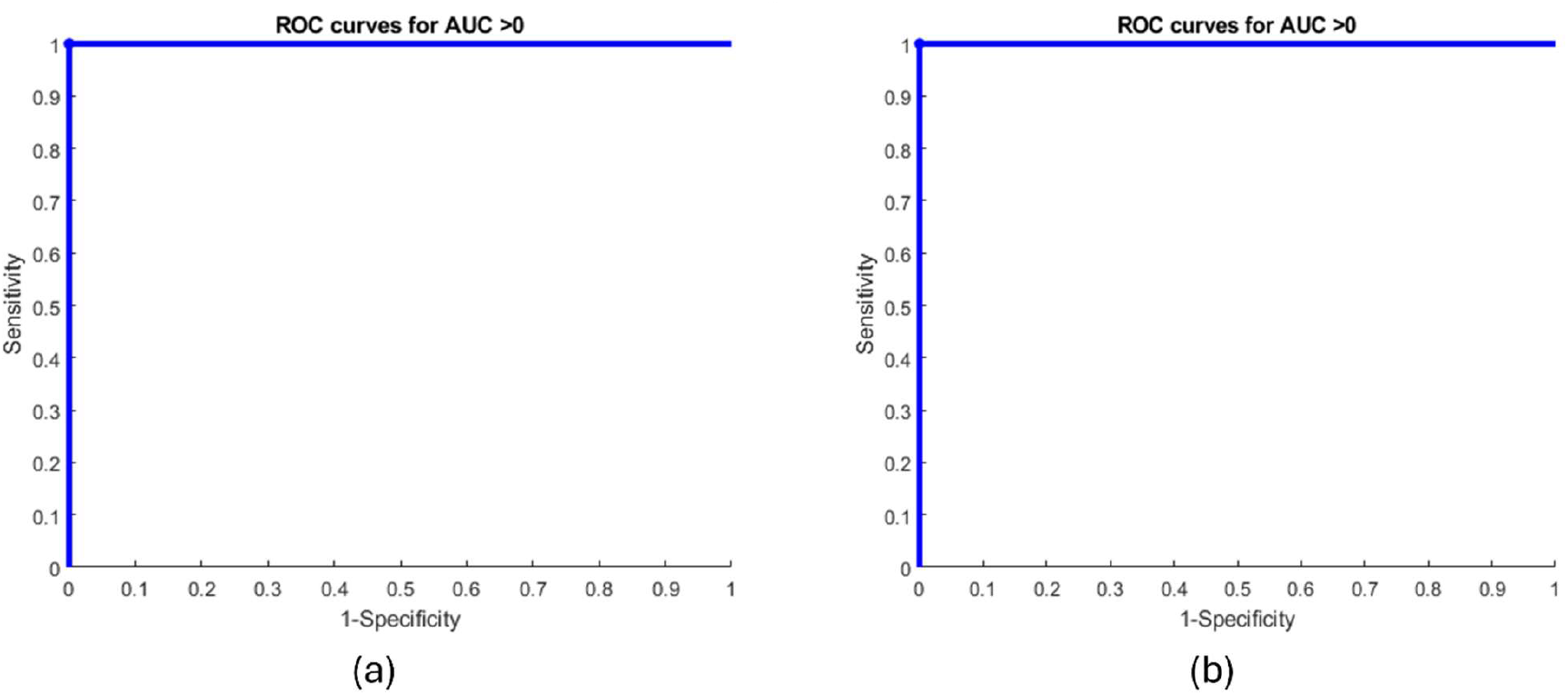
(a) ROC for dementia classifier performance for 100 bootstraps of 80:20 training:test split on in 105 plasma samples, where half are cases the other half are controls. Cases are identified through a combination of MRI scans, MMSE and CDR score determination; (b) ROC for gender classifier performance for 100 bootstraps of 80:20 training:test split on in the same plasma samples, where 60% are women and 40% are men. All bootstrap splits have the same proportion of men and women as in the complete dataset.

In addition to the phenotyping exercise described above, we used the broad-spectrum nano electrochemical vibrational sensor to quantitatively assay for molecular analytes in mixtures, demonstrating the concept and analysis workflow first with simple mixtures and then applying the approach to samples with increasing biochemical complexity of the background matrix, like plasma. Here, we describe the quantitative measurement of insulin glargine/lispro in mixtures of the two insulin isoforms, glargine (Humalog) and lispro (Toujeo), which are structurally different from one another in three amino acid units along their respective peptide chains [3]. Lispro is known to be a fast-acting isoform that enables meal-time control of blood glucose, whereas glargine provides all-day, long-acting control of blood sugar. The differing acting mechanisms of the two insulin isoforms necessitates that manufactured batches should be of free of cross contamination before they are administered to patients in clinical or in at-home settings. Multiple batches of mixtures of the two isoforms were prepared and serially diluted to test for detection sensitivity simultaneously as well. Concentration of the analyte isoforms varies between 0 ag/ml to 125 ag/ml in the batches, where 0 ag/ml concentration indicates a ‘mixture’ comprised of one isoform only. 2µl aliquots of the samples of different mixtures were pipetted into the electrochemical cell, scanned thrice per sensor over the -0.3V to 0.3 V bias range and the current-voltage-time measurements were post processed to remove contribution of aggressors to the electrochemical data as described before. Each batch was measured in triplicate with three different sensors with the tip-based instantiation of the sensor and all screened measurements are subsequently featurized to generate the electrochemical basis functions and their respective weights. An aligned UMAP reduction [8] of vibrational spectra of the two isoforms at 100ag/ml concentration is depicted in Figure 5a. To perform a conceptual quantitative assay for one isoform in a mixture, we project the high dimensional data from the mixture samples into a lower dimensional signal space that is most like the pure component samples containing the analyte isoform, and maximally different from the pure component samples that comprise of the second isoform. In addition, the signal space is also characterized by maximum contrast between samples corresponding to different mixture ratios, to enable discrimination between different stoichiometries of the isoforms in the reduced signal. These transforms are encapsulated into an eigenspace reduction of the scattering matrix, *S*, for a specific analyte isoform concentration dataset:

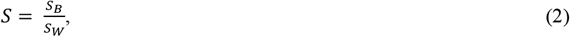

where *S*_*B*_ is the scattering matrix between the sample and pure non-analyte isoform data, as well as between the sample concentration aliquot data and other distinct mixture stoichiometries and *S*_*W*_ represents the scattering matrix between the sample and the pure analyte isoform datasets. The eigenspace reduction of *S* generates an optimally maximum representation of the scattering matrix wherein those sample features are down selected for which the sample is maximally distinct from the non-analyte isoforms and other stoichiometries while also most like the analyte isoform [9]. Analyte isoforms can be present at the same concentration when the non-analyte isoform can be mixed in at variable levels, and each stoichiometric ratio is treated as an independent sample dataset. Plots of the largest scattering matrix eigenvalue versus the concentration of the analyte isoform in the sample mixture are depicted in Figures 5b and 5c, where the analytes in 5b and 5c are glargine and lispro respectively. Each insulin isoform has an associated calibration curve as discovered from the eigenvalue decomposition. Error bars associated with each datapoint represent the spread in eigenvalues for different non-analyte isoform concentrations given a stoichiometric concentration value for the analyte isoform. The interpolated limit of detection for glargine and lispro as determined from the calibration curve are 0.1 and 2 ag/ml respectively, assuming the eigenvalue spread remains consistent about 0 ag/ml for each isoform type. This exercise demonstrates the ability to quantitate the amount of a specific analyte in a mixture that comprises of the analyte and a chemically similar molecular species, by generating a calibration curve that maps the reduced dimensionality signal to the concentration of the analyte in a functionally monotonic manner.

**Figure 5:**
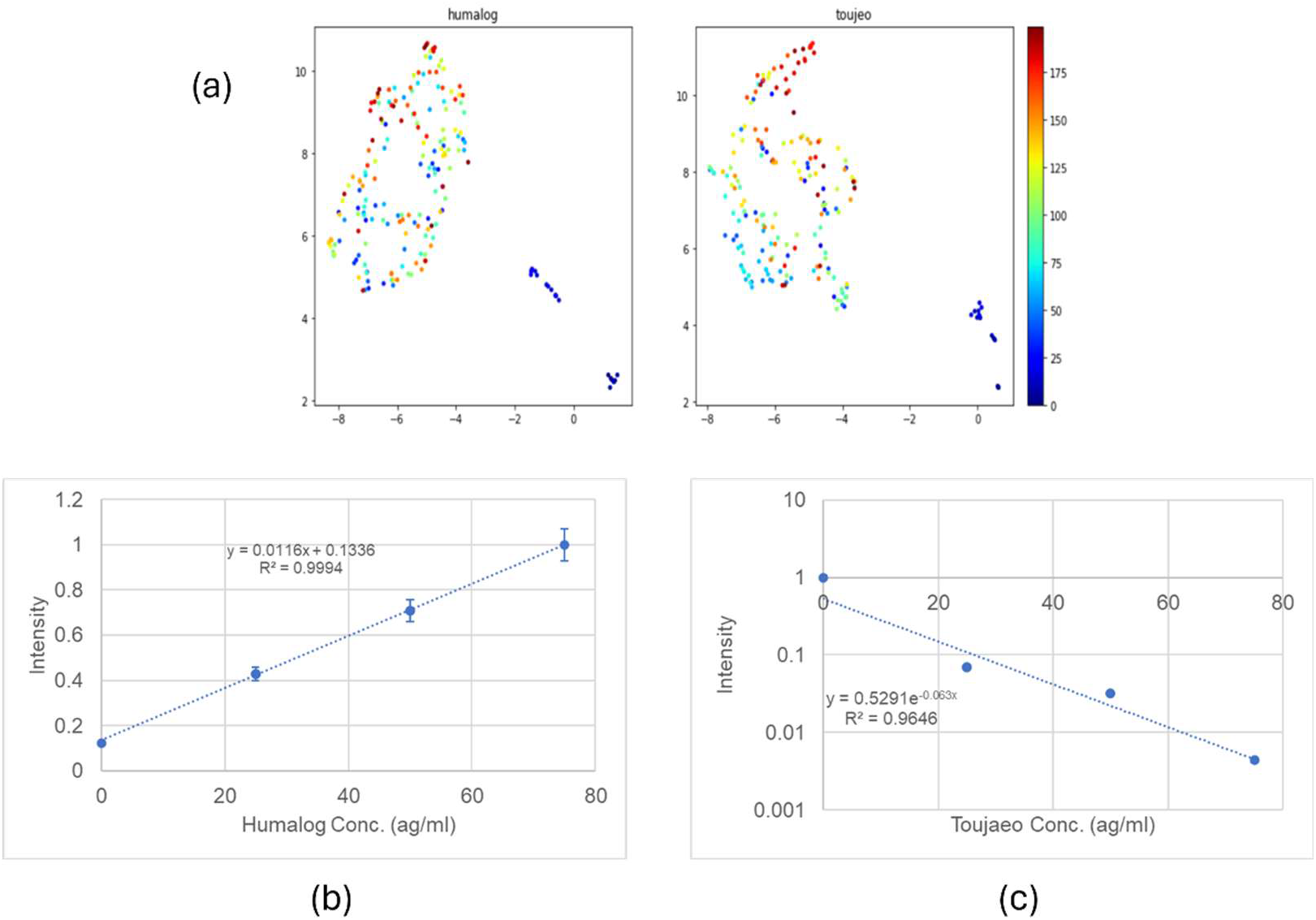
(a) aligned UMAP reduction for reference samples of Humalog (glargine) and Toujeo (lispro) at 100 ag/ml in the redox electrolyte, depicting the qualitative similarities and differences in the electrochemical vibrational spectra for the two peptide molecules; (b) calibration signal for Humalog in a batch of Toujeo; (c) calibration signal for Toujeo in a batch of Humalog.

We apply the analysis tools outlined above to also develop an assay for pro-inflammatory cytokine IL6 in a much more complex biological matrix, namely human plasma. A reference source of the protein was dissolved in PBS and used to prepare aliquots for spiking in the electrolyte buffer, as well as in the human plasma. The concentrations of the IL6 in the electrolyte as-prepared are 0, 0.025, 0.05, 0.5, 1, 2.5, 5, 10, 25, 50, 100, 500, 1000, 5000, 10000 pg/ml, and in the plasma matrix are 0, 0.025, 0.05, 0.1, 0.5, 1, 2.5, 5, 10, 25, 50, 100, 500, 1000 pg/ml. Each concentration aliquot in the two different matrices is measured in six technical replicates, after pipetting 4ul of the sample into the electrochemical cell, where the samples, diluted by the cell electrolyte, are scanned by the nano-electrochemical sensor over the -0.2V to 0.7 V bias range, with each measurement comprising of three scans per sensor. The scans were completed using vial-based reduction-to-practice for the sensor. The measurement data is screened for quality and transformed into the electronic energy-vibrational frequency feature space using the methods described previously. The feature files for electrolyte-only samples containing IL6, we created an estimate for the calibration signal of IL6 in the electrolyte matrix using the scattering matrix approach in Equation 2. Specifically, we identify the set of features in the feature matrix for a particular concentration aliquot that are a) most similar to other measurements of that aliquot, b) most similar to the highest concentration aliquots (5000, 10000 pg/ml) and c) most divergent from all other aliquots, with the scattering matrix ratio approach. In addition, the subspace of features identified from the scattering matrix approach are further filtered to identify the features that vary monotonically in intensity with concentration. The normalized norm of this reduced feature set is plotted against concentration of IL6 in the electrolyte matrix in Figure 6a. The calibration signal preserves linearity down to 5 pg/ml for the full dynamic range from 0 -1000 pg/ml. We also generate an equivalent calibration curve for a smaller dynamic range between 0-5 pg/ml, with significant linearity down to 0.025 pg/ml in Figure 6b. Next, we depict the application of an ensemble-based regression model, comprised of tree, random forest, SVM, kNN, linear, ridge and neural net regression models for the assay of IL6 in these plasma samples for the full range and for the smaller dynamic range at lower concentrations. The full measurement dataset for the model training and testing workflow comprises of the electrochemical features from samples comprised of IL6 spiked in the reference electrolyte and in plasma samples. For purposes of inferencing, specific concentration aliquots are held out from the complete measurement dataset and used for testing, where the remaining data is split 80:20 into training and validation data to train and optimize the model over. The features input to the model are pre-selected from the performance of a simple SVM model on a reduced set of features in the training data, where the features are apriori selected based on different criteria like minimal variance within cohorts or correlation strength between feature and label. The ensemble regularization parameter, the component model parameters and the ensemble weights are hyper-parameter optimized on the validation dataset. The results of the bootstrapped inferencing are depicted in Figure 6c, for the full dynamic range regressor, where each point is the median of all of the bootstrapped inferencing outcomes on the corresponding concentration aliquot. The error bars represent the inferencing variance for that specific concentration aliquot for all the trained models over the multiple bootstraps, which arises from the different training sets used in each of the bootstrap iterations. For this larger dynamic range, we see the lowest coefficient of variation in the predictions at ∼5%. Results of bootstrapped inferencing for the smaller dynamic range regression model at lower concentration values are shown in Figure 6d, with the lowest coefficient of variation being 18.5% at this reduced dynamic range. For context in Figures 6c and 6d, the red line is the y = x line that is indicative of perfect concordance between predicted and actual concentration values on the plasma samples. For both the large and small dynamic range models, the inferencing is conducted on 100 bootstrap splits of the training and testing data. As can be observed from the inferencing graph, the predictions show a high degree of correspondence with the actual concentration values.

**Figure 6:**
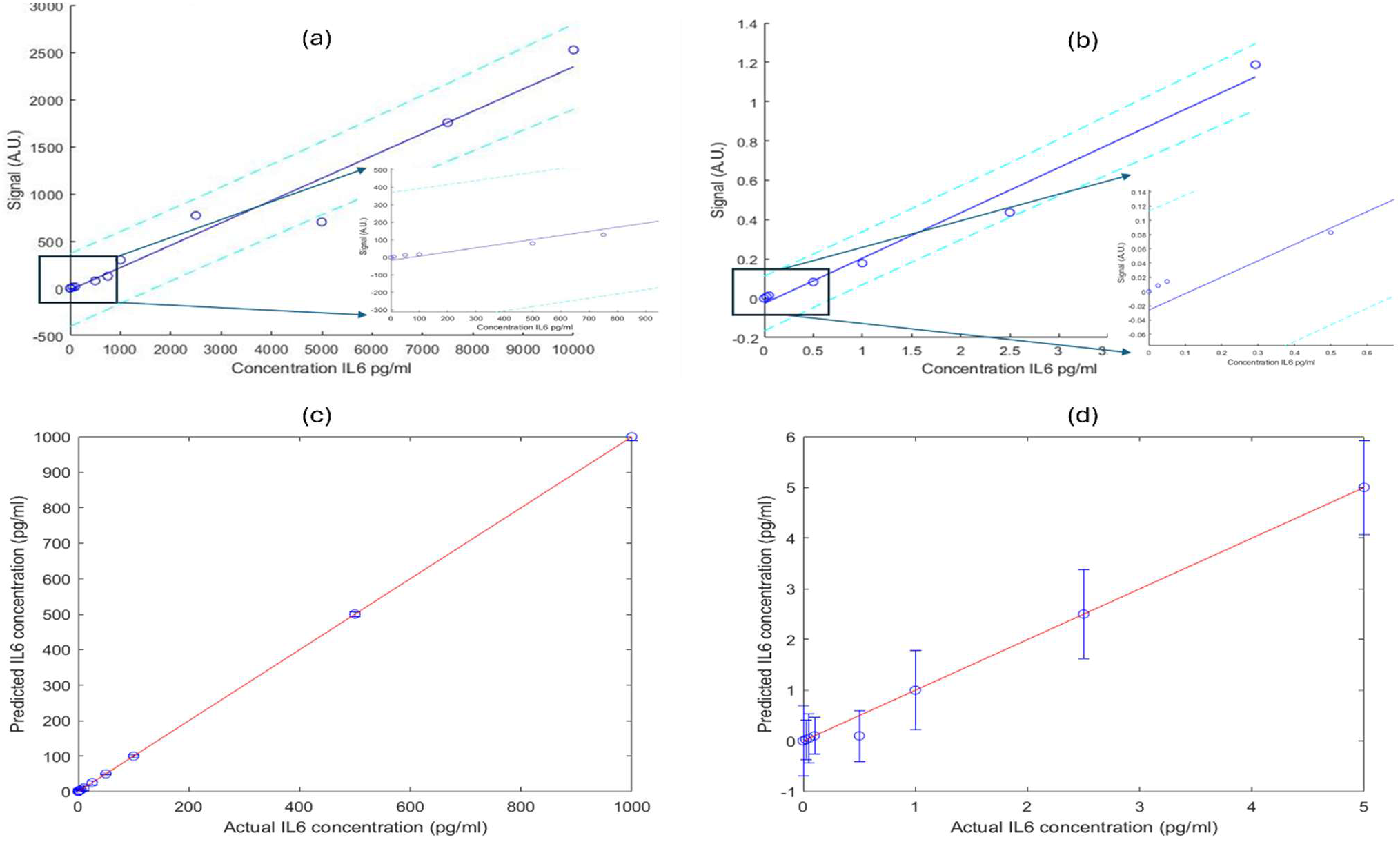
Calibration signal for IL6 for a dynamic range (a) from 0-10000 pg/ml and (b) from 0-5 pg/ml. Solid blue line represents the least square fit, and the dotted cyan lines represent the 2 S.D. spread around the least squares linear fit. The inference performance for models from (a) and (b) are depicted in (c) and (d). Performance is exhibited as the trendline between predicted concentration values on the y axis and the actual concentration values on the x axis, where the ideal performance is the y = x line, which is shown by the red dotted line. The smallest coefficient of variation is ∼ 5% on the full dynamic range in (c) and ∼ 18.5% for the reduced dynamic range model at low concentration values in (d). The model in (d) is triggered when concentration inferences in model (c) are below 10 pg/ml.

## Conclusions

We have demonstrated a novel analytical paradigm that combines nanoscale electrochemical sensing with machine learning to perform phenotype- and marker-agnostic quantitative and predictive assays of complex biological samples. The approach successfully liberates sample measurement and analysis from the constraints of hypothesis-driven, marker-specific probe and label methodologies by leveraging the broad-spectrum vibrational frequency transducing capabilities of a nanometer-scaled electrochemical interface, coupled with advanced feature extraction and machine learning algorithms.

The platform’s versatility was demonstrated across three distinct analytical challenges of increasing complexity. Most significantly, we demonstrated the platform’s ability to perform phenotypic classification by successfully distinguishing plasma samples from patients with dementia from age-matched controls with 100% sensitivity and specificity across several bootstrap iterations. We also demonstrated the ability of the platform to classify patient genders in a marker agnostic manner with 100% accuracy, using the same sample measurement dataset but different patient metadata, to create a new digital assay without requiring new physical measurements. This marker-agnostic approach to disease phenotyping represents a paradigm shift from traditional biomarker-based diagnostics toward pattern recognition of complex molecular signatures that may capture disease and other phenotypes more comprehensively than single analyte measurements. Future work will involve validation of these phenotypic in the clinical context with larger sample numbers, as well as investigating other disease phenotypes in biological samples like plasma, serum, whole blood, saliva, urine, sweat and CSF.

Subsequently, we achieved quantitative differentiation and measurement of structurally similar insulin isoforms (glargine and lispro) that differ by only three amino acid residues, with detection limits reaching 0.1 ag/ml for glargine and 2 ag/ml for lispro, in a simple one-component background matrix comprising of the other isoform. A scattering matrix-based approach was utilized to isolate the marker-specific calibration signal in the electrochemical feature matrix. This represents a significant advancement in the ability to detect and quantify minute concentrations of closely related biomolecules without requiring marker-specific reagents or preparation protocols.

Finally, we extended the methodology to assay IL-6 cytokine in human plasma, a considerably more complex biological matrix. The ensemble regression model achieved reliable quantitation, with a lower limit of quantitation at 25 fg/ml, demonstrating the platform’s capability to maintain analytical performance even in the presence of thousands of potentially interfering plasma proteins and metabolites. The linear response observed across the physiologically relevant concentration range (0.025 pg/ml to 1 ng/ml) validates the applicability of this approach for inflammatory biomarker detection. Future work will explore the inter- and intra-assay variability of the IL6 assay, as well as the dependence of the coefficient of variation on sample volume variability.

The analytical workflow’s foundation on electrochemical transduction of vibrational frequency information provides several key advantages over conventional analytical methods. The requirement of only 2-4 μl sample volumes addresses critical limitations in applications where sample availability is severely restricted. The unified measurement protocol eliminates the need for marker-specific sample preparation, reagents, or probes, potentially reducing cost, complexity, and time-to-result while increasing analytical flexibility.

The machine learning framework’s ability to extract meaningful signal from high-dimensional electrochemical feature spaces demonstrates the power of computational approaches in unlocking information content that would be inaccessible through traditional analytical methods. The ensemble modeling approach, incorporating multiple algorithmic perspectives (SVM, random forest, neural networks, k-nearest neighbors, etc.), provides robust predictions while minimizing overfitting risks through cross-validation, regularization and bootstrapping strategies.

From a broader perspective, this work represents a fundamental shift toward digitally-enabled analytical chemistry, where analytical capabilities are delivered through validated computational models rather than physical reagent kits. This paradigm has the potential to democratize advanced analytical capabilities by reducing infrastructure requirements and enabling rapid deployment of new assays through digital updates rather than reagent development cycles.

Several areas warrant further investigation to fully realize the platform’s potential. The scalability of the manufacturing process for the nanoscale electrochemical interfaces needs significant validation for commercial deployment and for clinical applications. Additionally, the robustness of the machine learning models across diverse sample matrices and population demographics requires extensive validation studies. The platform’s performance with other classes of analytes, including nucleic acids, lipids, and small molecule metabolites, will be systematically evaluated to define the full scope of analytical applications.

The platform’s ability to capture broad molecular signatures in a low-cost, unified workflow opens possibilities for discovering previously unknown biomarker patterns associated with disease states, potentially advancing and scaling both diagnostic and therapeutic development.

In conclusion, we have established a new analytical framework that combines the molecular specificity of nanoscale electrochemical interfaces with the pattern recognition capabilities of machine learning to enable phenotype- and marker-agnostic analysis of complex biological samples. The demonstrated applications in protein quantitation and disease phenotyping validate the platform’s potential to address current limitations in biological sample analysis and point toward a future where analytical chemistry is increasingly defined by computational models rather than chemical reagents.

## References

(1) Clark, K. D., Zhang, C., Anderson, J. L., “Sample Preparation for Bioanalytical and Pharmaceutical Analysis”, Anal. Chem., 88, 11262 (2016)

(2) Gupta, C., Walker, R. M., Chang, S., Fischer, S. R., Seal, M., Murmann, B., Howe, R. T., “Quantum Tunneling Currents in a Nanoengineered Electrochemical System”, J. Phys. Chem. C, 121, 15085 (2017)

(3) Donnor, T., Sarkar, S., “Insulin-Pharmacology, Therapeutic Regimens and Principles of Intensive Insulin Therapy”, Endotext (https://www.endotext.org), South Dartmouth (MA), MDText.com, Inc., 2000-

(4) Gupta, C., Pena-Perez, A., Fischer, S. R., Weinreich, S. B., Murmann, B., Howe, R. T., “Active control of probability amplitudes in a mesoscale system via feedback-induced suppression of dissipation and noise”, J. App. Phys., 120, 224902 (2016)

(5) Jones, S. A., Takeuchi, T., Aletaha, D., Smolen, J., Choy, E. H., McInnes, I., “Interleukin 6: The biology behind the therapy”, Consid. Med., 2, 2 (2018)

(6) Friedman, J. H., Popescu, B. E., “Predictive learning via rule ensembles”, Ann. App. Stat., 2, 916 (2008)

(7) Dong, X., Yu, Z., Cao, W., Gan, L., Shi, Y., “A survey on ensemble learning.” Trans. Art. Int., 1, 10 (2020)

(8) Dadu, A., Satone, V. K., Kaur, R., Koretsky, M. J., Iwaki, H., Qi, Y. A., Ramos, D. M., Avants, B., Hesterman, J., Gunn, R., Cookson, M. R., Ward, M. E., Singleton, A. B., Campbell, R. H., Nalls, M. A., Faghri, F., “Application of Aligned-UMAP to longitudinal biomedical studies”, Patterns, 4, 100741 (2023)

(9) Ghifary, M., Balduzzi, D., Kleijn, W. B., Zhang, M., “Scatter Component Analysis: A Unified Framework for Domain Adaptation and Domain Generalization”, IEEE Trans. Patt. Anal. Mach. Learning, 39, 1414 (2017)

(10) Bard, A.J., and Faulkner, L.R., “Electrochemical Methods: Fundamentals and Applications”, John Wiley & Sons, 2^nd^ edition, 2001

(11) Hastie T., Tibshirani R., and Friedman J., “The Elements of Statistical Learning: Data Mining, Inference, and Prediction”, Springer, 2^nd^ edition, 2009

(12) Belkin, M., Hsu, D., Ma, S., Mandal, S., “Reconciling modern machine-learning practice and the classical bias–variance trade-off”, Proc. Nat. Acad. Sc., 116, 15849 (2019)

