## Supplemental Information for "Machine Learning-Augmented Analysis of Nano-electrochemical Sensor Data for Predictive and Quantitative Assays of Complex Biological Samples"

The process of feature generation/extraction from the raw current-voltage-time datasets is described in this section. An electrochemical measurement of the sample in the electrochemical cell generates matrices of measured current versus time and measured voltage versus time per scan, where rows correspond to the measured variable at a given applied bias and columns correspond to the measured variable at a given time point in the data acquisition cycle. These observation variables qualify the electrostatic state of the nano-electrochemical interface across which the vibrationally assisted-electronic transition occurs. Here, we present an approach to map these observational variables to generalized parameters that define the vibrational-electronic energy exchange mechanism that underpins the charge transfer process.

Equation 5.4 of [2] describes a first order rate constant for the rate of charge transfer that is of an "intermediate" nature between adiabatic and non-adiabatic processes, from one vibronic energy level in the donor species to another vibronic energy level in the acceptor species. The rate constant is a normalized representation of the observed transition rate when integrated over the plurality of full donor and empty acceptor states. The theoretical transition rate, in this integral form, is a superposition of multiple vibrational frequency dependent terms, each of which is representative of the probabilistic transition path for the transferring electrons when assisted with a specific vibrational mode. Additional events that occur before and after the vibrationally-activated transition that are thermally activated and electrostatics-independent are wrapped into path dependent constants $w_i$ that multiply into the vibrational-electronic transition probability as seen below:

$$I \propto \sum_i \int_{E_e} \int_{E_{\Omega_i}} w_i P\left(E_e - E_{\Omega_i}, \Omega_i, [\Theta]\right) f_e(E_e) \left(1 - f_{\Omega_i}\left(E_{\Omega_i}\right)\right) \quad \text{(S1)}$$

Where $E_e, E_{\Omega_i}$ represent the energy levels of the donor and acceptor states, $f_e(E_e), f_{\Omega_i}\left(E_{\Omega_i}\right)$ are the occupancy probabilities of those energy levels in the donor and acceptor states and $P\left(E_e - E_{\Omega_i}, \Omega_i, [\Theta]\right)$ is the probability of the vibrationally-assisted electronic transition for the mode $\Omega_i$ that depends on the applied bias, $E_e - E_{\Omega_i}$ and other parameters $[\Theta]$ characterizing the electrostatic environment around the charge transfer interface, as defined in Equation 5.4 of [2]. For purposes of calculating the integral, and for the nano-electrochemical transducer described in the paper, $f_e(E_e)$ is the Fermi-Dirac distribution probability and $f_{\Omega_i}\left(E_{\Omega_i}\right)$ is the Gaussian redox occupation probability in the electrolyte [10]. From (1), the transition current is now represented as a superposition of a finite number ($i = 1 \ to \ n$) of basis terms that have closed form parametrized and analytic expressions. Alternatively, the transition current can be thought of as an output from a shallow, single layer neural network, comprised of $i$ parametrized neurons, weighted by $w_i$, given a set of applied bias values $E_e - E_{\Omega_i}$ and environmental parameters $[\Theta]$, with a loss function that minimizes the regularized least squares error between the observed and predicted $I$ values. We determine the parameter and weight/path dependent probability values using a stochastic gradient with momentum solver approach for optimization on the LSE loss function, where 80% of the matched current-voltage values randomly sampled from the measurement matrices are used as training data, and the shallow neural network is validated on the remaining 20% held-out data. This training/validation split is shuffled every epoch of the model learning, the learning rate is dropped in a piecewise manner every 10 epochs, and each training iteration occurs over a mini batch of samples to estimate the gradient of the loss function and to evaluate the path probabilities. The set of parameters and path-probabilities constitute a new set of features that the original current-voltage measurements are reduced into for subsequent analysis. Note, this reduction occurs for a specific, fixed value of number of neurons/basis terms for signal reduction.

The superposition representation of the transition flux observables in S1 allows us to analytically estimate the expected bias and variance of this transformation, following the approach by [11]:

$$Bias \sim (1 - W^T(WW^T + (M^T)^{-1}II^TM^{-1})^{-1}W), \quad \text{(S2a)}$$

$$Var \sim W^T(WW^T + (M^T)^{-1}II^TM^{-1})^{-1}W(W^T(WW^T + (M^T)^{-1}II^TM^{-1})^{-1}W)^T, \quad \text{(S2b)}$$

Where,

$$M = \int_{E_e} \int_{E_{\Omega_i}} P\left(E_e - E_{\Omega_i}, \Omega_i, [\Theta]\right) f_e(E_e) \left(1 - f_{\Omega_i}\left(E_{\Omega_i}\right)\right) \quad \text{(S2c)}$$

$W$ is the matrix of path-dependent probabilities ($[w_1(\Theta)\ w_2(\Theta) \ldots w_N(\Theta)]$) for different environmental parameter values, $I$ is the observed transition current and $E_e - \langle E_{\Omega_i} \rangle_\Omega$ are the measured bias values. The global parameter optimization routine described above is used to estimate $W$ and $[\theta]$, from which the bias and variance values can be estimated. These bias and variance estimates, are determined as functions of the number of vibrationally assisted channels $n$, are combined into a risk metric and depicted in Figure S-1. The double descent risk profile [12] demonstrates an interpolation threshold at $n$ = 165, and models with higher $n$ continue to interpolate the training data with high fidelity while also exhibiting low generalization risk in testing. To ensure feature transformation within 30 minutes, we select $n$ = 300.

Figure S1

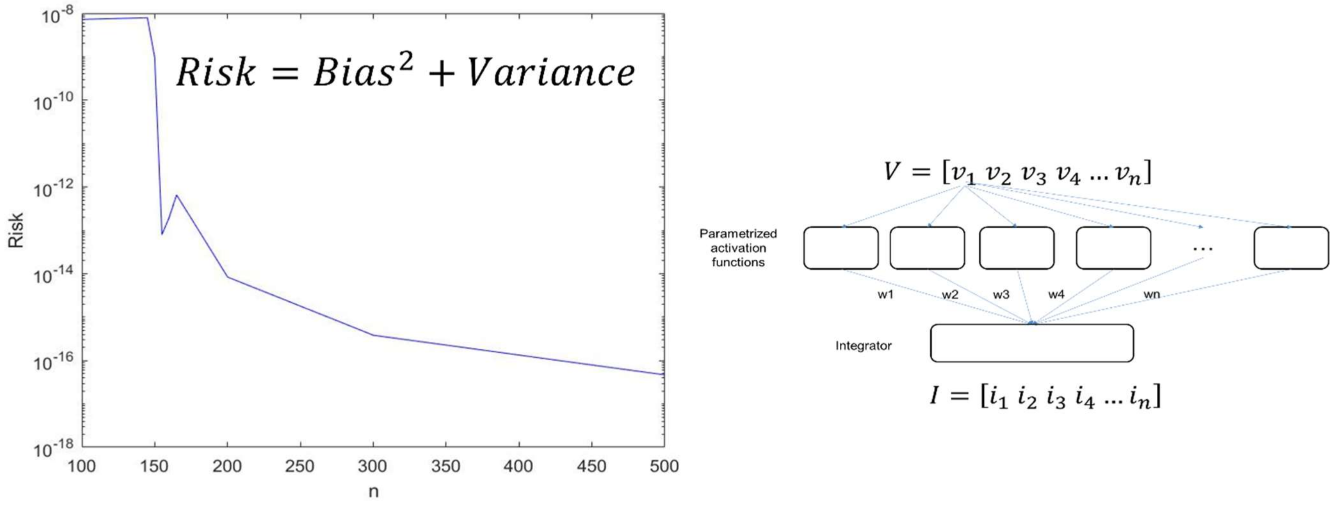


Figure S1: Model prediction risk versus model complexity, as denoted by feature dimensionality. Interpolation region is at n = 165. The risk depicts the double descent function that comprises of under parametrization to the left of interpolation region and over parametrization to the right of the interpolation region. The value of n determines the size of the shallow neural network model that is used to determine the vibrational features from the observed current and voltage data.
